# Cancer-like fragmentomic characteristics of somatic variants in cell-free DNA

**DOI:** 10.1101/2025.04.28.651130

**Authors:** Zhenyu Zhang, Yunyun An, Mengqi Yang, Yuqi Pan, Xiaoyi Liu, Fanglei Gong, Huizhen Lin, Bianbian Tang, Wanqiu Wang, Yunxia Bai, Xin Zhao, Yu Zhao, Changzheng Du, Kun Sun

## Abstract

Cell-free DNA (cfDNA) in plasma consists of short DNA fragments resulting from a non-random fragmentation process, with distinct “fragmentomic” characteristics that are related with their cellular origins. Despite the wealth of low-pass whole genome sequencing data from recent cancer liquid biopsy studies, somatic variants within this data have often been overlooked without detailed characterization. Here, we reported that in low-pass sequencing data, somatic variant signatures in cfDNA markedly differ between non-cancerous controls and cancer patients, indicating that tumor-associated signals are retained in these variants. Surprisingly, our investigation into cfDNA fragmentomics showed that even in controls, cfDNA molecules harboring somatic variants exhibit cancer-like traits, such as reduced size, decreased DNA methylation, and altered end motif usages and distributions in the nucleosome structure. Importantly, these somatic variants-associated fragmentomic aberrations are more pronounced in cancer patients, enabling cancer diagnosis. In a large pan-cancer cohort, we utilized artificial intelligence to integrate genomic, fragmentomic, and epigenomic features, developing diagnostic models named FreeSV (<u>Fr</u>agmentomic and <u>e</u>pigenomic <u>e</u>xaminations of <u>S</u>omatic <u>V</u>ariants in cfDNA) and FreeSV-m. Leveraging somatic variant-associated features alone, the FreeSV model achieved area under the ROC curves (AUCs) between 0.81-0.92 across cancer types; however, when genomewide features were also included, the AUCs of FreeSV-m model substantially increased to 0.93-0.99 across cancer types, highlighting the significance of integrative genomic and fragmentomic analyses in cfDNA for cancer liquid biopsy.

## Introduction

Circulating cell-free DNA (cfDNA) molecules in human plasma are small fragments of DNA that originate from dead cells (e.g., through apoptosis or necrosis)^1^. In non-cancerous subjects, cfDNA is primarily sourced from the hematopoietic system and liver^2–4^; in contrast, in cancer patients, the tumors release cfDNA into plasma pool^5^, which serves as a valuable analyte for non-invasive diagnostics, a process commonly referred to as “liquid biopsy”^6^. Tumor-derived cfDNA not only carry somatic mutations but also retain epigenomic modifications such as DNA methylation^7,8^. Furthermore, recent studies indicate that the fragmentation patterns, termed “fragmentomics”, of tumor-derived cfDNA differ from those of non-tumor-derived cfDNA. These patterns are influenced by endonuclease activities and preferences, including shorter fragment sizes^9^, altered end or breakpoint motif usages^10,11^, and increased fractions of ends cut within nucleosomes^12^, all of which have gained substantial attention as emerging biomarkers in cancer liquid biopsy^8,13–15^.

In most cancer patients, tumor-derived cfDNA usually account for a small proportion of the total cfDNA pool in plasma, posing significant challenges for accurately identifying tumor-derived somatic mutations. To overcome this limitation, one widely utilized strategy is to sequence cfDNA at extra-high depth, either through whole genome sequencing or targeted capture^16–18^. For instance, in our previous study involving a hepatocellular carcinoma (HCC) patient, we sequenced cfDNA to approximately 220x human haploid genome coverage, alongside genotyping paired white blood cells to profile and filter out germline variants^18^. Recent advancements in computational algorithms, such as cfSNV, necessitate at least 200x coverage to detect mutations present at a 5% frequency in cfDNA^19^. In contrast, the majority of current liquid biopsy studies, particularly those focusing on cfDNA fragmentomics^8,13^, typically generate data with less than 5x human haploid genome coverage, often lacking information on germline genotypes. Additionally, various factors like germline polymorphisms, technical artifacts (including PCR and sequencing errors), aging, and clonal hematopoiesis introduce variants into cfDNA^20–23^, making it nearly impossible to accurately identify tumor-derived somatic mutations from low-pass sequencing data.

Interestingly, large-scale cancer studies, such as The Cancer Genome Atlas (TCGA), have revealed that somatic mutations in tumors exhibit non-random patterns, known as “mutational signatures”. The COSMIC (Catalogue of Somatic Mutations in Cancer) database has catalogued a curated list of mutational signatures in pan-cancer, along with annotations pertaining to cancer types and associated pathologies^24,25^. A recent investigation by Wan et al. demonstrated that the mutational signatures in cfDNA from cancer patients differ from those of controls, thereby showing potential in cancer diagnosis^26^. Additionally, Bruhm et al. reported genome-wide alterations in mutation frequencies and types among cancer patients^27^, underscoring the merit of analyzing somatic variants within low-pass cfDNA datasets. Given the substantial volume of low-pass cfDNA data generated in cancer liquid biopsy studies, including those generated via whole genome bisulfite sequencing^28^, Enzymatic Methyl-seq (EM-seq)^29^ or similar protocols, analyzing somatic variants might yield novel biomarkers that enhance diagnostic performance and provide insights into cfDNA biology, particularly regarding fragmentomic characteristics linked to somatic variants that remain unexplored. In this study, we examine the characteristics of somatic variants in low-pass cfDNA without paired germline genotypes. Beyond mutational signatures, we further investigate the cfDNA fragmentomics and epigenomics associated with variants and assess their potential in constructing cancer diagnostic models.

## Results

### Schematic workflow for genomic and fragmentomic characterizations of somatic variants

For each low-pass cfDNA dataset comprising both control and cancer samples, we identified somatic variants in the absence of germline genotyping information. A stringent filtering strategy was employed, utilizing data quality metrics and known polymorphism sites (see Methods). We then profiled and compared the genomic mutational signatures between controls and cancer samples to elucidate tumor-associated footprints within the somatic variants, focusing specifically on single-base substitutions (SBSs). Subsequently, we extracted cfDNA molecules encompassing the somatic variants and examined their fragmentomic and epigenomic characteristics (for EM-seq data only) in comparison to those covering reference alleles of the same loci. Lastly, we leveraged artificial intelligence (AI) to integrate the genomic, fragmentomic, and epigenomic features of cfDNA somatic variants, facilitating the development of cancer diagnostic models aimed at exploring the translational value of somatic variants in cfDNA.

### Genomic and fragmentomic profiles of somatic variants in low-pass cfDNA data

We first examined a cohort of hepatocellular carcinoma (HCC) samples from our previous study, which included 56 HCC samples and 24 age- and gender-matched non-cancerous controls^12^. All cfDNA samples in this cohort were sequenced to ∼3-5x human haploid genome coverage (Suppl. Fig. S1a). Following stringent filtering, we obtained 100,000-150,000 somatic variants (i.e., candidate mutations) for each sample. Notably, while there were no statistical differences in sequencing depths between HCC and control samples, the number of somatic variants were significantly higher in HCC samples compared to controls at equivalent sequencing depths (P=1.7×10^-4^, Chow test; Suppl. Fig. S1b-c). We then calculated frequencies of the mutation types according to sequence context for each sample. The results indicated that mutation profiles were generally similar between HCC and control samples, with increased frequencies in C>T and T>C transitions (Fig. 1a; Suppl. Table S1). However, Principal Component Analysis (PCA) and unsupervised clustering both revealed systematic differences between HCC samples and controls (Fig. 1b-c). As depicted in Fig. 1c, the samples clustered into two groups: 35 out of 40 samples were HCC in Cluster-1, which was significantly higher than Cluster-2 where only 21 out of 40 samples were HCC (P=0.0015, Chi-Squared test). Furthermore, the top-ranked mutation contexts significantly elevated in the HCC-enriched Cluster-1, including C[C>T]G, G[C>T]G, C[T>C]G, and G[T>C]G (all P<10^-5^, Mann-Whitney U tests with Benjamini & Hochberg adjustment), have been previously reported as common mutation types in HCC and various cancers^25,30^.

**Fig. 1.**
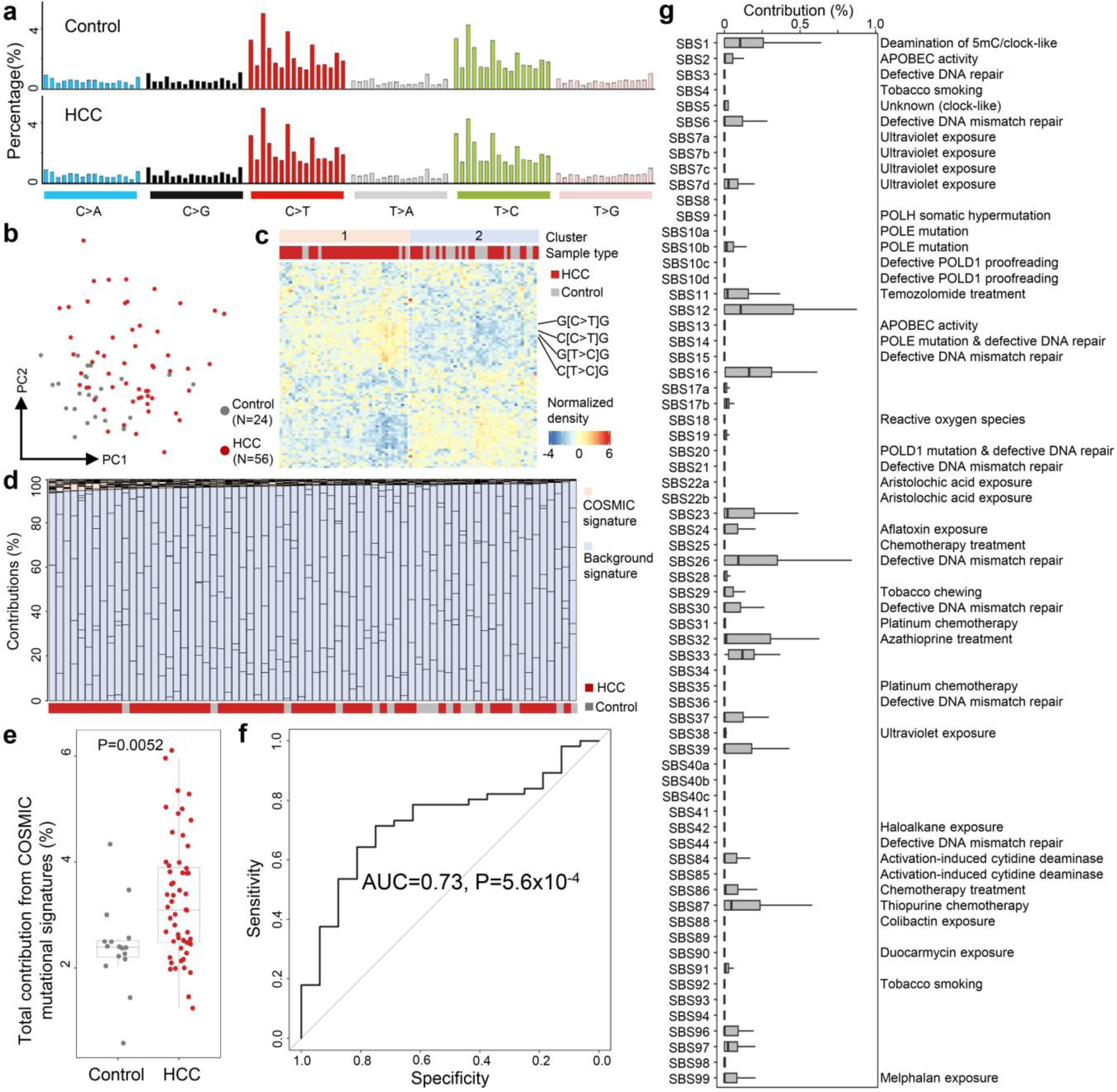
Genomic profiles of somatic variants in cfDNA in our Hepatocellular Carcinoma (HCC) cohort. **a)** Mutation profiles of a non-cancerous control and an HCC sample. **b)** Principal Component Analysis and **c)** unsupervised clustering results based on mutation profiles. **d)** Deconvolution results against a pool of COSMIC mutational signatures and “background signatures” from controls. **e)** Total contributions from COSMIC mutational signatures between controls and HCC samples after deconvolution. P-value was calculated using Mann-Whitney U test. **f)** Receiver Operating Characteristic curve analysis for differentiating HCC patients from controls using contributions from COSMIC mutation signatures after deconvolution. P-value was calculated using Z-test. **g)** Contributions of COSMIC mutational signatures in HCC samples. In **b,e**, each dot represents one sample. In **e,g**, boxplots represent the median, upper and lower quartiles and whiskers indicate 1.5x IQR.

To further investigate the biological implications of the mutation profiles, we deconvoluted them against known tumor mutational signatures. Given that tumor-derived DNA constituted only a small fraction of the cfDNA pool, besides the 67 mutational signatures for single-base substitutions documented in the COSMIC database (excluding signatures annotated as “sequencing artifacts”), we also randomly selected 8 control samples to model the mutation patterns typical of non-cancerous backgrounds, which may arise from technical artifacts and clonal hematopoiesis. For all samples, high cosine similarities (>0.999 in all samples; Suppl. Fig. S2a) were achieved between the original mutation profiles and the fitting results, confirming successful deconvolutions^26^. As expected, both HCC and control samples predominantly reflected contributions from the “background signatures” (Fig. 1d). However, HCC samples exhibited significantly higher contributions from COSMIC mutational signatures compared to controls (P=0.0052, Mann-Whitney U test; Fig. 1e). Receiver operating characteristic (ROC) curve analysis indicated that the total contributions of COSMIC mutational signatures effectively differentiated HCC samples from controls, with an area under the ROC curve (AUC) value of 0.73 (P=5.6×10^-4^, Z-test; Fig. 1f). Additionally, the total contributions of COSMIC mutational signatures displayed a positive correlation with the tumor DNA fractions in HCC samples (P=6.8×10^-4^, Spearman correlation; Suppl. Fig. S2b). Of the 67 COSMIC mutational signatures, SBS1, SBS12, SBS16, SBS26, and SBS33 showed notable contributions in HCC samples (Fig. 1g), with SBS1, SBS12, and SBS16 previously recognized as commonly present in HCC^25^. Collectively, these findings suggest that the mutation profiles in low-depth cfDNA samples contain substantive tumor-associated information beyond mere background noises, warranting further investigations.

### Fragmentomic features of cfDNA harboring somatic variants

Next, we examined the fragmentomic characteristics associated with the somatic variants identified in our HCC cohort. For all loci with somatic variants, we extracted cfDNA reads harboring the variant alleles (referred to as “Mut-DNA” hereafter) alongside those covering reference alleles (referred to as “Wt-DNA”). We observed that, for both controls and HCC samples, Mut-DNA fragments were shorter than Wt-DNA, as indicated by significantly higher fractions of short fragments (i.e., ≤ 150 bp; both P<10^-10^, paired t-tests; Fig. 2a-c). We quantified the difference in fractions of short fragments between Mut- and Wt-DNA for each sample, termed “Diff-size”. As shown in Fig. 2d, Diff-size values were significantly greater in HCC samples than in controls (P=0.012, Mann-Whitney U test), with an AUC value of 0.68 for differentiating the two groups (P=0.0033, Z-test; Fig. 2e). We then investigated two well-studied motif features: the 5’-CCCA end motif and the CT-5’-CC breakpoint motif (Fig. 2f-k)^10,11,31^. The 5’-CCCA end motif exhibited a significant reduction in Mut-DNA among controls, while no significant differences were found for the CT-5’-CC breakpoint motif (P=0.025 and 0.70, respectively, paired t-tests). As contrast, both motifs were significantly underrepresented in Mut-DNA from HCC samples (both P<0.001; paired t-tests). To quantify the differences in motif usage between Mut- and Wt-DNA, we defined “Diff-CCCA” and “Diff-CTCC”. We found a significant difference in Diff-CTCC between HCC and controls, while it was not statistically different for Diff-CCCA (P=0.043 and 0.19, respectively, Mann-Whitney U tests). Moreover, Diff-CTCC exhibited an AUC value of 0.64 for distinguishing HCC samples from controls (P=0.020, Z-test; Fig. 2e).

**Fig. 2.**
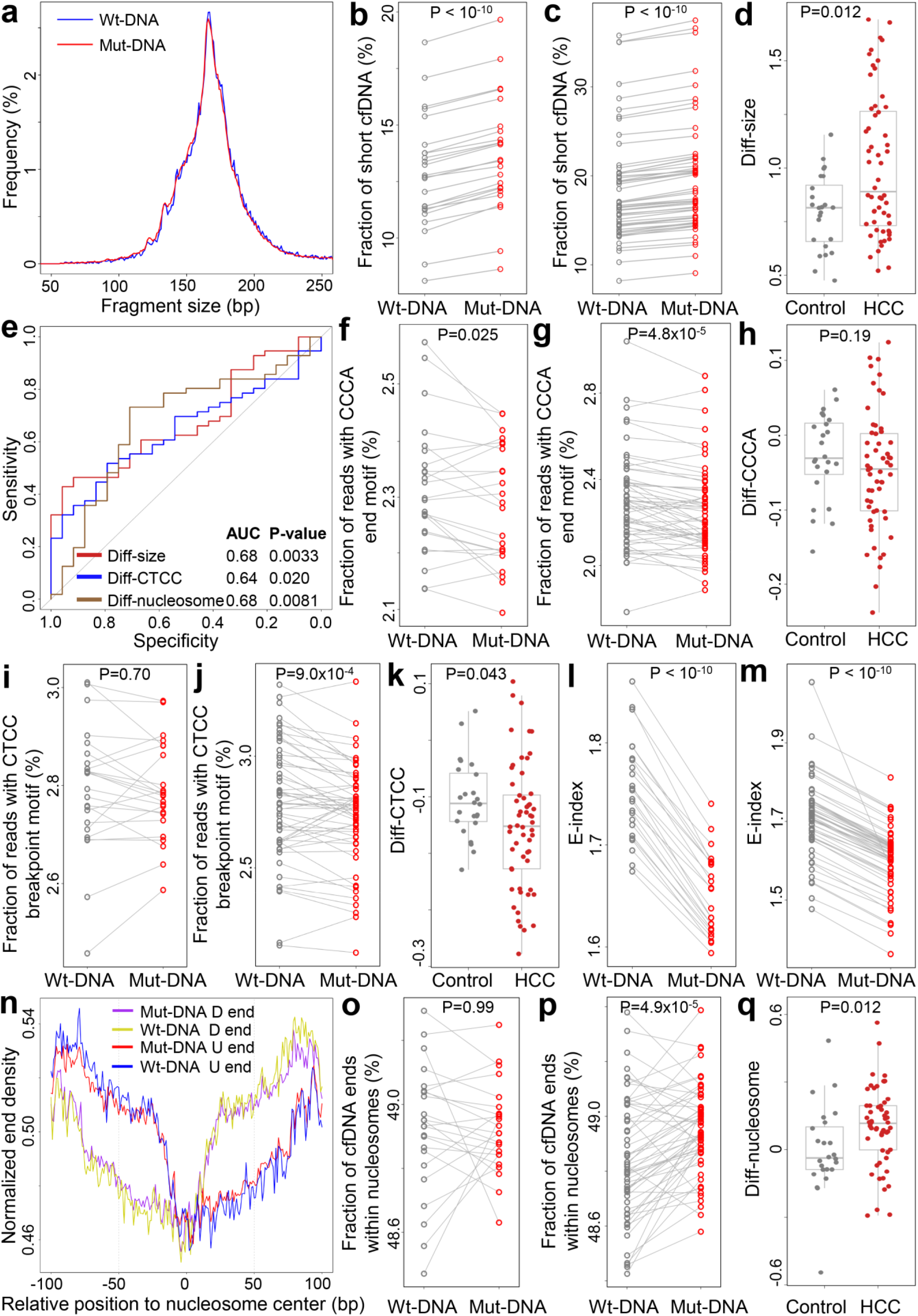
Fragmentomic features of cfDNA associated with somatic variants in the HCC cohort. **a)** Size distributions of cfDNA compassing variant (Mut-DNA) and reference (Wt-DNA) alleles in the variant loci in an HCC sample. **b-c)** Fraction of short cfDNA fragments (defined as shorter of equal to 150 bp) between Wt- and Mut-DNA in **b)** control subjects and **c)** HCC patients. **d)** Differences in fractions of short fragments between Mut- and Wt-DNA (Diff-size) between controls and HCC samples. **e)** ROC curves for differentiating HCC patients from controls using various fragmentomic features. **f-g)** Fraction of reads with 5’-CCCA end motif for Wt- and Mut-DNA in **f)** controls and **g)** HCC samples. **h)** Differences in fraction of reads with 5’-CCCA end motif (Diff-CCCA) between Mut- and Wt-DNA in controls and HCC samples. **i-j)** Fraction of reads with CT-5’-CC breakpoint motif for Wt- and Mut-DNA in **i)** controls and **j)** HCC samples. **k)** Differences in fraction of reads with CT-5’-CC breakpoint motif (Diff-CTCC) between Mut- and Wt-DNA in controls and HCC samples. **l-m)** E-index values for Wt- and Mut-DNA in **l)** controls and **m)** HCC samples. **n)** Orientation-aware fragment end distributions in the nucleosomal context in an HCC sample. U and D end stand for Upstream and Downstream end, respectively. **o-p)** Fraction of ends located within nucleosomes (defined as ±50 bp from the nucleosome center) for Wt- and Mut-DNA in **o)** controls and **p)** HCC samples. **q)** Differences in fraction of ends located within nucleosomes (Diff-nucleosome) between Mut- and Wt-DNA in controls and HCC samples. In **b,c,f,g,i,j,l,m,o,p**, P-values were calculated using paired t-tests. In **d,h,k,q**, P-values were calculated using Mann-Whitney U test. In **e**, P-values were calculated using Z-tests. In **b-d,f-m,o-q**, each dot represent one sample. In **d,h,k,q**, boxplots represent the median, upper and lower quartiles and whiskers indicate 1.5x IQR.

Lastly, we investigated the distribution of fragment ends in both Mut- and Wt-DNA. Notably, the E-index values, a metric assessing the consistency of fragment ends in a given sample compared to a panel of healthy controls^1^, were significantly lower for Mut-DNA than for Wt-DNA in both control and HCC samples (both P<10^-10^, paired t-tests; Fig. 2l-m). This finding indicates a greater discordance between the ends of Mut-DNA and those from healthy subjects. However, there was no statistically significant difference in the extent of E-index decrease between these two groups (Suppl. Fig. S3). We further profiled the end distributions of Mut- and Wt-DNA within the nucleosomal context of GM12878 cells (lymphoid lineage), a commonly used nucleosome track in cfDNA fragmentomic analyses, in an orientation-aware manner^1,12,32–34^. As shown in Fig. 2n-p, Mut-DNA exhibited a higher fraction of ends located within nucleosomes (defined as ±50 bp from the nucleosome center) compared to Wt-DNA in HCC samples (P=4.9×10^-5^, paired t-tests), but such difference was not observed in controls. Additionally, we quantified the difference in the fraction of ends located within the nucleosomes between Mut- and Wt-DNA, referred to as “Diff-nucleosome”. This analysis revealed significant differences between HCC samples and controls (P=0.012, Mann-Whitney U test; Fig. 2q), and Diff-nucleosome demonstrated an AUC value of 0.68 for differentiating the two groups (P=0.0081, Z-test; Fig. 2e). Collectively, these results suggest that cfDNA encompassing somatic variants exhibit cancer-like fragmentomic features, even in non-cancerous individuals. Furthermore, the pronounced differences in fragmentomic characteristics were more evident in HCC samples, indicating their potential as biomarkers for cancer diagnosis.

### Somatic variant-associated cfDNA fragmentomics in mouse model

To validate the cancer-like fragmentomic features of somatic variants in control subjects, we collected and analyzed cfDNA samples from C57BL/6J strain mice (N=12). As these mice are genetically identical to the reference genome of this strain^35^, all detected somatic variants in cfDNA were attributed to clonal hematopoiesis or technical artifacts rather than germline polymorphisms. We identified somatic variants in these samples and extracted cfDNA molecules covering somatic variants (i.e., Mut-DNA) as well as those covering reference alleles at the same loci (i.e., Wt-DNA), following the approach used in human data analysis. The size distribution of Mut- and Wt-DNA from a representative murine cfDNA sample was illustrated in Fig. 3a. Consistent with the human data, Mut-DNA showed significantly shorter size compared to Wt-DNA (P=5.0×10^-7^, paired t-test; Fig. 3b). Moreover, we did not observe significant differences in 5’-CCCA end motif usage between Mut- and Wt-DNA (P=0.70, paired t-test; Fig. 3c); however, Mut-DNA exhibited a significant reduction in CT-5’-CC breakpoint motif usage compared to Wt-DNA (P=0.0036, paired t-test; Fig. 3d). Collectively, these findings conformed alterations in cfDNA fragmentomics associated with Mut-DNA, indicating that clonal hematopoiesis might influence cfDNA fragmentation patterns.

**Fig. 3.**
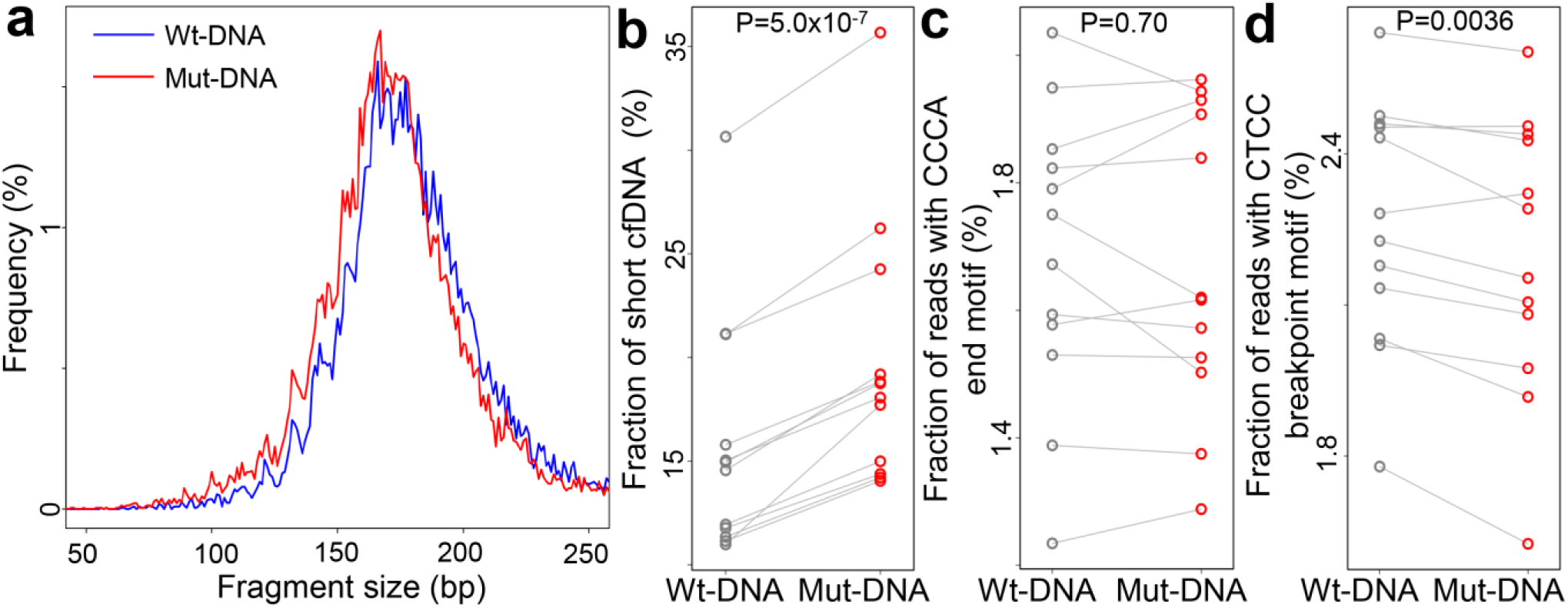
Somatic variant-associated cfDNA fragmentomics in murine samples (N=12). **a)** Size distributions of cfDNA compassing variant (Mut-DNA) and reference (Wt-DNA) alleles in one sample. Comparisons in fractions of **b)** short cfDNA fragments, **c)** reads with 5’-CCCA end motif, and **d)** reads with CT-5’-CC breakpoint motif between Wt-DNA and Mut-DNA. In **b-d**, P-values were calculated using paired t-tests.

### Fragmentomic features of somatic variants in cfDNA in other cancer types

To validate our findings, we analyzed a public cfDNA dataset from Liang et al., which includes samples from 10 hepatocellular carcinoma (HCC) patients, 8 lung cancer patients, and 10 healthy controls^36^. The mutation patterns identified in this dataset were consistent with those observed in our HCC cohort, with a clear preference for C>T and T>C variants (Fig. 4a and Suppl. Table S2). PCA and unsupervised clustering of the mutational signatures revealed a distinct separation between control and cancer samples (Fig. 4b and Suppl. Fig. S4). We further used all 10 control samples as background references to deconvolute the mutational signatures in the cancer samples. As shown in Fig. 4c, the majority of mutational contributions in both HCC and lung cancer samples were attributed to the “background signatures”, which corroborates the findings in our HCC cohort (Fig. 1d). Notably, several COSMIC mutational signatures highly represented in our HCC cohort, SBS1, SBS16, SBS26, and SBS33, also showed prominent contributions in the HCC samples of this dataset. In contrast, lung cancer samples exhibited increased contributions from SBS30 and SBS44 compared to HCC samples, while contributions from SBS26 were reduced (Fig. 4d). These results suggest that cfDNA compassing somatic variants preserve tumor-specific information across different cancer types, with distinct mutational signatures observed between HCC and lung cancer.

**Fig. 4.**
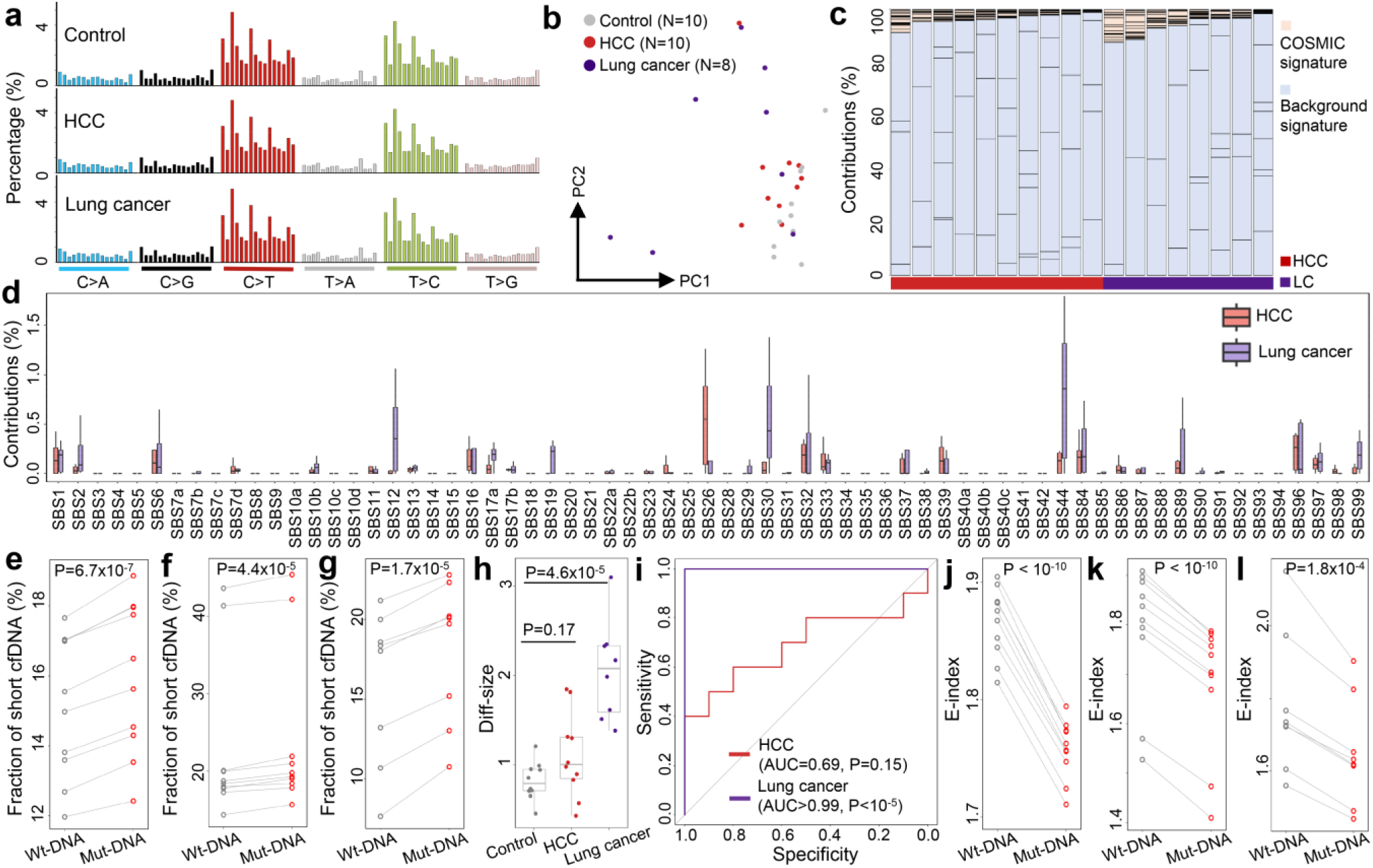
Genomic and fragmentomic features of somatic variants in cfDNA in Liang et al. cohort. **a)** Mutation profiles of a non-cancerous control, an HCC sample, and a lung cancer sample. **b)** PCA result based on the mutation profiles. **c)** Deconvolution results, and **d)** contributions of COSMIC mutational signatures after deconvolution. **e-g)** Fraction of short cfDNA fragments for Wt- and Mut-DNA in **e)** controls, **f)** HCC samples, and **g)** lung cancer samples. **h)** Diff-size values among controls and cancer samples. **i)** ROC curves for differentiating cancer patients from controls using Diff-size values. **j-l)** E-index values for Wt- and Mut-DNA in **j)** controls, **k)** HCC samples, and **l)** lung cancer samples. In **e,f,g,j,k,l**, P-values were calculated using paired t-tests. In **h**, P-values were calculated using Mann-Whitney U tests. In **i**, P-values were calculated using Z-tests. In **b,e-h,j-l**, each dot represent one sample. In **d,h**, boxplots represent the median, upper and lower quartiles and whiskers indicate 1.5x IQR.

Next, we extracted Mut- and Wt-DNA from each sample and investigated their fragmentomic characteristics. The results are summarized in Fig. 4e-l and Suppl. Fig. S5. Briefly, Mut-DNA was consistently shorter than Wt-DNA across all sample groups with statistically significant differences (P < 10^-4^ for all groups, paired t-tests; Fig. 4e-g). Additionally, Diff-size values were significantly higher in lung cancer samples, enabling perfect differentiation from controls (Fig. 4h-i). While Diff-size also showed a similar trend in HCC samples, the difference did not reach statistical significance. Furthermore, E-index values for Mut-DNA were significantly lower than for Wt-DNA across all sample groups (Fig. 4j-l). Together, these results further substantiate our findings from the HCC cohort, showing that Mut-DNA exhibits cancer-associated fragmentomic features in both cancer patients and non-cancerous controls.

### Fragmentomic features of variants in Enzymatic Methyl Sequencing of cfDNA

Considering that DNA methylation in cfDNA has been extensively studied in cancer liquid biopsy, to explore whether we could profile somatic variants and fragmentomics in such data, a pan-cancer EM-seq dataset from Bie et al. was collected and analyzed^37^. After quality control, we retained 1,185 cfDNA samples, including 448 non-cancerous controls and 737 cancer samples from seven types: 64 breast cancer (BRCA), 135 colon/rectal cancer (COREAD), 56 esophageal cancer (ESCA), 109 gastric cancer (STAD), 107 liver cancer (LIHC), 150 non-small cell lung cancer (NSCLC), and 116 pancreatic cancer (PACA) for further analysis. First, we identified somatic variants and mutation profiles across all samples. Overall, the mutation profiles were similar to those observed in whole genome sequencing (Fig. S6 and Suppl. Table S3). PCA of the mutation profiles revealed that control samples clustered together, while cancer samples exhibited substantial heterogeneity (Fig. 5a). Unsupervised clustering identified 2 major clusters with consistent trend: 412 out of 448 controls (92.0%) were in Cluster-1, significantly higher than random assignment (P=3.3×10^-4^, binomial test). Moreover, 56.5% samples in Cluster-1 were cancerous, while cancerous samples accounted for 84.9% in Cluster-2 (P < 10^-10^, Chi-Squared test; Fig. 5b), suggesting distinct mutation profiles between cancer and control samples.

**Fig. 5.**
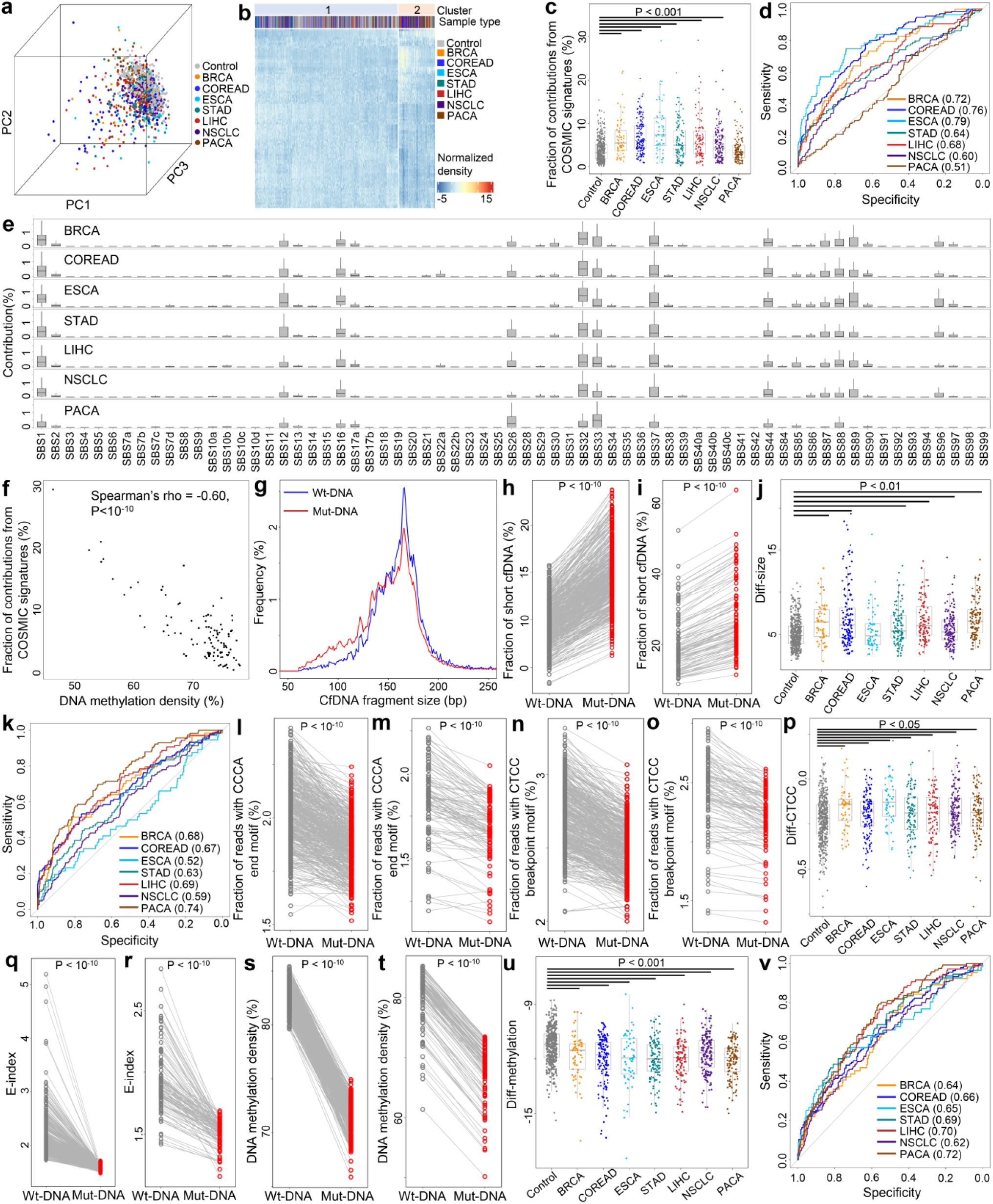
Genomic, fragmentomic, and epigenomic features of somatic variants in cfDNA in Bie et al. dataset. **a)** PCA and **b)** unsupervised clustering results using mutation profiles. **c)** Total contributions of COSMIC mutational signatures across controls and cancer samples of different types. **d)** ROC curves for differentiating cancer samples from controls using total contributions from COSMIC mutational signatures. The AUCs were recorded in parentheses. All P<0.05 except PACA calculated by Z-tests. **e)** Contributions of COSMIC mutational signatures across different cancer types. **f)** Correlation between total contributions of COSMIC mutational signatures and overall DNA methylation levels in liver cancer samples. **g)** Size distributions of Wt- and Mut-DNA in a liver cancer sample. **h-i)** Fraction of short cfDNA fragments in Wt- and Mut-DNA in **h)** controls and **i)** liver cancer samples. **j)** Diff-size values across controls and cancer samples. **k)** ROC curves for differentiating cancer samples from controls using Diff-size. The AUCs were recorded in parentheses. All P<0.05 except ESCA calculated by Z-tests. **l-m)** Fraction of reads with 5’-CCCA end motif for Wt- and Mut-DNA in **l)** controls and **m)** liver cancer samples. **n-o)** Fraction of reads with CT-5’-CC breakpoint motif in Wt- and Mut-DNA in **n)** control subjects and **o)** liver cancer patients. **p)** Diff-CTCC values across controls and cancer samples. **q-r)** E-index values in Wt- and Mut-DNA in **q)** controls and **r)** liver cancer samples. **s-t)** DNA Methylation levels in Wt- and Mut-DNA in **s)** controls and **t)** liver cancer samples. **u)** Differences in methylation levels (Diff-methylation) between Mut- and Wt-DNA across controls and cancer samples for CpGs within gene bodies, and **v)** corresponding ROC curves for differentiating cancer samples and controls. AUCs were recorded in parentheses, and all P<0.001 calculated by Z-tests. In **h,i,l-o,q-t**, P-values were calculated using paired t-tests. In **c,j,p**, P-values were calculated using Mann-Whitney U tests. In **d,k**, P-values were calculated using Z-tests. In **a,c,f,h-j,l-u**, each dot represent one sample. In **c,e,j,p,u**, boxplots represent the median, upper and lower quartiles and whiskers indicate 1.5x IQR. BRCA: breast cancer, COREAD: colon/rectal cancer, ESCA: esophageal cancer, STAD: gastric cancer, LIHC: liver cancer, NSCLC: non-small cell lung cancer, PACA: pancreatic cancer.

Next, we randomly selected 10 control samples as background references and deconvoluted the mutation profiles of the remaining samples against 67 COSMIC mutational signatures and the background signatures. For all samples, the majority of mutational contributions were attributed to the “background signatures”; however, the contributions from COSMIC mutational signatures were significantly higher in cancer samples from 6 cancer types compared to controls (all P < 0.001, except PACA, Mann-Whitney U tests; Fig. 5c). ROC analysis revealed AUC values ranging from 0.60 to 0.79 for differentiating these 6 cancer types from controls (all P < 0.001, Z-tests; Fig. 5d). Common pan-cancer mutational signatures, such as SBS1, SBS12, SBS16, SBS32, SBS33, and SBS37, were shared across cancer types, while cancer-specific signatures, such as SBS85 and SBS90, were also identified (Fig. 5e). Moreover, the total contributions of COSMIC mutational signatures were negatively correlated with overall DNA methylation levels in all cancer types (all P < 10^-4^, Spearman correlation; Fig. 5f and Suppl. Fig. S7). As tumor-derived cfDNA molecules are hypomethylated ^7^, which leads to a negative correlation between overall DNA methylation levels and tumor DNA fractions in cfDNA data^38^, the results suggest a positive correlation between the total contributions of COSMIC mutational signatures and tumor DNA fractions, echoing the results in our HCC cohort (Suppl. Fig. S2b).

We next examined the fragmentomic features associated with somatic variants by extracting Mut-DNA and Wt-DNA from each sample. As shown in Fig. 5g-i and Suppl. Fig. S8, Mut-DNA was again significantly shorter than Wt-DNA in both cancer and control samples (P < 10^-10^ for all groups, paired t-tests). Diff-size values were significantly higher in 6 cancer types compared to controls (all P < 0.01, except ESCA, Mann-Whitney U tests; Fig. 5j), with AUC values ranging from 0.59 to 0.74 for differentiating these cancer samples from control samples (all P < 0.01, Z-tests; Fig. 5k). Additionally, the use of 5’-CCCA end motif and CT-5’-CC breakpoint motif were significantly reduced in Mut-DNA compared to Wt-DNA in both control and cancer samples (all P < 10^-10^, paired t-tests; Fig. 5l-o and Suppl. Fig. S9). Notably, Diff-CTCC values were significantly elevated in all cancer types compared to controls (all P < 0.05, Mann-Whitney U tests; Fig. 5p), while Diff-CCCA values did not differ significantly in most cancer types (Suppl. Fig. S9g). Furthermore, Mut-DNA exhibited significantly lower E-index values than Wt-DNA across all groups (all P < 10^-10^, paired t-tests; Fig. 5q-r and Suppl. S10), while no significant differences were observed in the fraction of reads located within nucleosomes between Mut-DNA and Wt-DNA (Suppl. Fig. S10). Together, the results confirm the aberrant fragmentomic features associated with somatic mutations in cfDNA, consistent with findings from whole genome sequencing datasets.

As the dataset was generated using EM-seq protocol, we also analyzed DNA methylation profiles associated with somatic variants. Considering that DNA methylation is context-dependent^39^, we narrowed the analysis to CpG sites covered by both Mut- and Wt-DNA; in addition, we divided the CpG sites into 3 groups based on their genomic contexts: promoters (defined as ± 500 bp around transcription start sites), gene bodies, and intergenic regions. As a result, in all samples (including controls), Mut-DNA exhibited significantly lower DNA methylation levels than Wt-DNA for all 3 groups of CpGs (P < 10^-8^ for all groups, paired t-tests; Fig. 5s-t and Suppl. Fig. S11-S12). We then quantified the differences in methylation levels between Mut- and Wt-DNA for each sample, referred to as “Diff-methylation”. As shown in Fig. 5u, Diff-methylation values for CpGs within gene bodies were significantly lower in all 7 cancer types compared to controls (all P < 0.001, Mann-Whitney U tests; Fig. 5u and Suppl. Fig. S11); ROC analysis demonstrated that Diff-methylation values for CpGs within gene bodies could differentiate cancer samples from controls with AUCs ranging from 0.62 to 0.72 (all P < 0.001, Z-tests; Fig. 5v). In contrast, Diff-methylation values for CpGs in promoters did not show significant differences between cancer samples and controls, and Diff-methylation values for CpGs in intergenic regions were significantly lower in 3 cancer samples compared to controls (Suppl. Fig. S12). These findings suggest that, in addition to fragmentomic features, epigenomic alterations also existed in cfDNA harboring somatic variants.

### Artificial Intelligence-empowered cancer diagnostic models

Building on the findings presented in Fig. 5, we applied artificial intelligence (AI) to integrate genomic, fragmentomic, and epigenomic characteristics associated with somatic variants in cfDNA to develop cancer diagnostic models on the Bie et al. dataset. The features included mutation profiles, Diff-size, Diff-CCCA, Diff-CTCC, Diff-nucleosome, and Diff-methylation for CpGs in gene bodies, totaling 101 features. We employed a gradient boosting machine (GBM) model and conducted 10-fold nested cross-validation repeated 100 times, using the averaged prediction scores to assess the cancerous status of the samples, as validated in previous studies^12,40^. We termed this approach FreeSV (Fragmentomic and Epigenomic Examination of Somatic Variants in cfDNA). FreeSV demonstrated AUC values ranging from 0.81 to 0.92 across the 7 cancer types, and 0.77 to 0.91 among different stages (all P < 10^-10^, Z-tests; Fig. 6a-b and Suppl. Fig. S13).

**Fig. 6.**
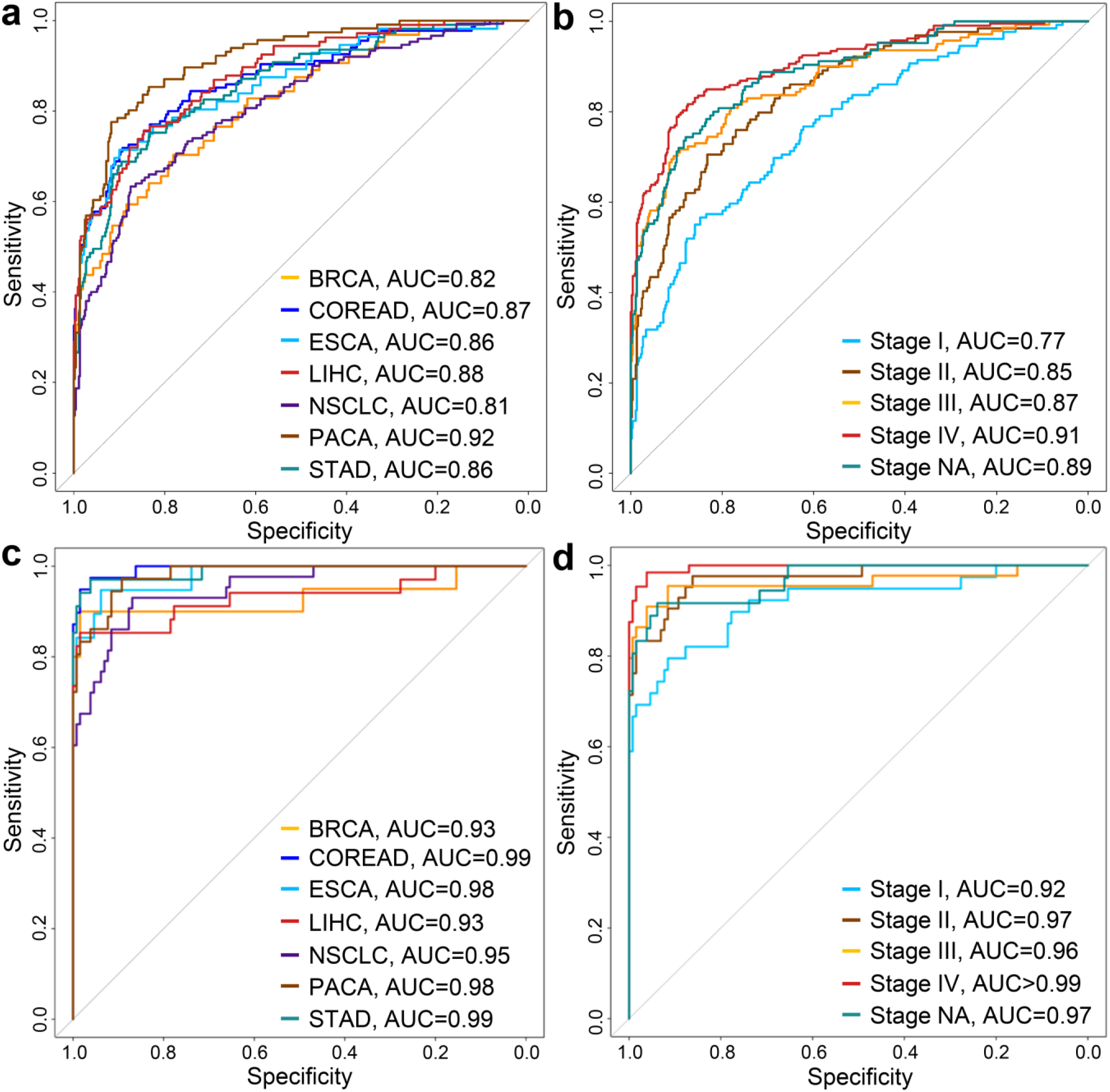
Performance of FreeSV and FreeSV-m models for cancer diagnoses. **a-b)** ROC curves of FreeSV model leveraging only somatic variants-associated features across **a)** cancer types and **b)** stages. **c-d)** ROC curves of FreeSV-m model leveraging both somatic variants-associated features and genomewide features on samples in the testing group across **c)** cancer types and **d)** stages. P<10^-10^ for all AUCs, Z-tests.

In practice, multiple features could be integrated towards building high-performance diagnostic models. For instance, in Bie et al. study, the authors incorporated 4 genomewide features (i.e., DNA methylation, fragment size, end motif, and copy number) and built a highly accurate model named THEMIS^37^. As a proof-of-concept, we combined our somatic variant-associated features with these 4 genomewide features and built a new model termed “FreeSV-m” to explore whether integration of our features could further improve the performance. For a fair comparison, as in Bie et al. study, the samples were divided into training and testing groups, and we built the model on the training group then directly applied the model on the testing group. On the training group, we measured the performance of FreeSV-m using 10-fold nested cross-validation repeated 100 times, which resulted in ROC curves with AUCs ranging from 0.96 to 0.99, all of which were higher than THEMIS (Suppl. Fig. S14). On the testing group, the AUCs were comparable between FreeSV-m and THEMIS (Fig. 6c-d); however, at 99% specificity, FreeSV-m correctly identified 183 out of 225 cancer samples (sensitivity=81.3%, 95% CI: 75.6-86.2%; Suppl. Table S4), which was significantly higher than THEMIS’s 74.4% (P=0.0076, binomial test); moreover, FreeSV-m correctly identified 71.6% stage I/II cancer samples (Suppl. Table S4), which was also significantly higher than THEMIS’s 61.0% (P=0.037, binomial test). Together, these results highlight the feasibility of integrating multi-omics features towards high-performance diagnostic models, as well as the translational significance of our somatic variant-associated features.

## Discussion

In this study, we performed an integrative analysis on the fragmentomic characteristics of somatic variants in cfDNA. Over the past decade, research focus in cancer liquid biopsy has partially shifted from genomic and epigenomic to fragmentomic characteristics of cfDNA^8,13,15^, which hold the advantage of working with shallow sequencing data. However, as technical artifacts and clonal hematopoiesis could introduce variants in cfDNA, analyzing genuine tumor-derived somatic mutations in low-pass cfDNA data remains highly challenging. Indeed, most of the variants found in cfDNA are not tumor-derived, but rather represent background noises, which can dilute the true tumor-associated signals in cfDNA from cancer patients. However, despite this problem, principal component analysis (PCA) and unsupervised clustering analyses both showed some degree of separation between control and cancer samples (Fig. 1b-c, 4b, and 5a-b), suggesting that even in low-pass cfDNA, tumor-associated signals can still be discerned from the somatic variants present in cancer patients. Overall, our findings on the genomic profiles of somatic variants in cfDNA are highly consistent with previous work by Wan et al.^26^, while our approach differs from that of Wan et al. in several key aspects. The primary difference is our use of a panel of control samples to simulate the background noises in addition to the COSMIC mutational signatures during mutation profile deconvolutions. As shown in Fig. 1d-e, 4c, and 5c, after deconvolution, the majority of contributions were actually attributed to the “background signatures”, while cancer samples showed higher contributions from COSMIC mutational signatures and the total contributions of COSMIC mutational signatures were positively correlated with tumor DNA fractions in cfDNA (Fig. 5f, Suppl. Fig. S2b, and S7). This suggests that our approach can effectively filter out noises, which are often related to experimental and analytical biases that are highly protocol/platform-dependent^41^, and emphasize the cancer-associated signals. Moreover, our approach was validated in both whole genome sequencing and EM-seq datasets, suggesting the robustness and generality of our approach.

Beyond genomic analysis, we conducted a comprehensive investigation on the fragmentomic characteristics in cfDNA associated with somatic variants, an area not previously explored in detail. Surprisingly, even in non-cancerous controls, cfDNA molecules covering somatic variants (“Mut-DNA”) exhibited cancer-like fragmentomic patterns compared to those covering reference alleles (“Wt-DNA”). These patterns included shorter fragment sizes, aberrant end or breakpoint motif usage, and higher proportions of ends located within nucleosomes. In a prior study, we demonstrated that DNA methylation plays a key role in cfDNA fragmentation, with lower methylation levels correlating with shorter cfDNA fragments^1^. Previous studies have also shown that mutations in DNMT3A and TET2, which occur during clonal hematopoiesis^42,43^, lead to decreased DNA methylation in aged hematopoietic cells^44,45^. Accordingly, Mut-DNA in the EM-seq data exhibited decreased DNA methylation levels (Fig. 5s-t), suggesting that in non-cancerous subjects, the age-related clonal hematopoiesis could be a major contributor of somatic variants in cfDNA and serves as the cause of cancer-like fragmentomic patterns in Mut-DNA. This finding thus provided evidences to elucidate the shortness of cfDNA molecules observed in aged people^1,46^ and highlighted the necessity of age-matched controls in cancer diagnosis^47^. Moreover, in cancer cfDNA samples, size patterns had been utilized as a valuable filter for the identification of tumor-derived mutations^18,48^; however, our findings suggested that such filter could not eliminate the clonal hematopoiesis-related variants as they are also associated with shorter cfDNA, which partially explained the false positives in previous studies^18,48^. In clinical, clonal hematopoiesis is associated with worse clinical outcome in various diseases, including cardiovascular diseases and cancer^49,50^; hence, our findings suggested the possibility of investigating clonal hematopoiesis by incorporating mutational profiles and fragmentomics in cfDNA^51^. Besides clonal hematopoiesis, a recent study reported cancer-like fragmentomic patterns in cfDNA from non-cancerous patients with TP53 mutations. In these cases, mutant TP53 altered chromatin accessibility, contributing to the observed fragmentomic changes^52^. Besides human data, somatic variant-associated cancer-like fragmentomic patterns (especially the relative shortness) was validated in murine cfDNA samples (Fig. 3). Together, the results indicated that investigating cfDNA fragments covering somatic variants in non-cancerous subjects offers novel insights into the regulation of cfDNA fragmentomics, shedding light on the mechanisms underlying cfDNA fragmentation and its implications for cancer diagnostics.

In addition, we defined several parameters to quantitatively assess the fragmentomic and epigenomic alterations between Mut- and Wt-DNA, including Diff-size, Diff-CCCA, Diff-CTCC, Diff-nucleosome, and Diff-methylation. Notably, in most analyses, these parameters revealed significant differences between cancer patients and controls. For example, across all datasets analyzed, Diff-size values were consistently higher in cancer patients than in controls, in line with the fact that tumor-derived cfDNA molecules are shorter than non-tumor-derived ones^1,9^. When examining breakpoint motif usage, Diff-CTCC values were lower in cancer patients in the HCC cohort, consistent with prior studies reporting reduced usage of the CT-5’-CC breakpoint motif in cfDNA from cancer patients^11^. Interestingly, in Bie et al. EM-seq dataset, Diff-CTCC values were elevated in cancer patients (Fig. 5p), which may be attributed to high biases in EM-seq data caused by cytosine-to-thymine conversions^53^. Further investigation is warranted to validate the reproducibility of this observation. Overall, these parameters hold promise as a novel category of biomarkers for cancer diagnosis. Furthermore, we developed FreeSV, an AI-enhanced model that integrates genomic, fragmentomic, and epigenomic features associated with somatic mutations in cfDNA. The validation of FreeSV using Bie et al. dataset with more than 1,100 samples demonstrates the translational merit of somatic variant analysis and the innovative fragmentomic features derived in this study, particularly when applied to low-pass cfDNA data. Moreover, as a proof-of-concept, we further integrated these somatic variant-associated features identified in this study with genomewide features leveraged in Bie et al. study. As shown in Fig. 6c-d and Suppl. Fig. S14-S15, the performance (especially the sensitivity) was significantly improved, which was better than using either somatic variants-associated features or genomewide features alone. The results thus demonstrated the translational significance of integrating multi-omics features in cfDNA towards building high-performance cancer diagnostic models.

In summary, this study provides a comprehensive profiling of genomic and fragmentomic characteristics in low-pass cfDNA data without paired germline genotypes. We show that cfDNA molecules harboring somatic variants in controls exhibit cancer-like fragmentomics compared to those covering reference alleles. Additionally, our computational approach allows for the extraction of novel biomarkers from low-pass cfDNA data at minimal cost, with potential for integration into high-accuracy cancer diagnostic models, especially when combined with other biomarkers through AI.

## Methods

### Ethics approval and sample processing

This study was approved by the Ethics Committee of Shenzhen Bay Laboratory. Animal study was conducted according to protocols approved by SZBL Animal Center. Wildtype C57BL/6J strain mice (N=12) were housed under specific pathogen-free conditions with a 12h light/dark cycle, at a temperature of 20-26°C, and a relative humidity of 40-70%; mice were fed a standard mouse chow diet and were sacrificed between 12-30 weeks. For both human subjects and mice, blood samples were centrifuged at 1,600 g, 4 °C for 15 min, and then the plasma portion was harvested and re-recentrifuged at 16,000 g, 4 °C for 15 min to remove blood cells^1,54^; plasma samples were stored at −80 °C until further usage. CfDNA extraction and library preparation were performed as in our previous study^12^. Briefly, for each mouse sample, cfDNA was extracted from 300-600 μL plasma with MagPure Circulating DNA LQ Kit (Magen, #IVD5432) and 3-6 ng cfDNA was used to construct libraries using VAHTS Universal DNA Library Prep Kit for MGI (Vazyme, #NDM607) following the manufacturers’ instructions. CfDNA libraries were sequenced on an MGISEQ-T7 (MGI) sequencer in paired-end 100 bp mode.

### Sequencing data processing

For whole-genome sequencing data, raw reads were first preprocessed using Ktrim software (version 1.5.0)^55^ to remove sequencing adapters and low-quality cycles. The preprocessed reads were then aligned to the reference human genome (NCBI GRCh38) or reference mouse genome of C57BL/6J strain “B6Eve” (Jackson Laboratory)^35^ for human and mouse data, respectively, using BWA-MEM software (version 0.7.17)^56^. Alignment results were sorted and indexed using SAMtools (version 1.17)^57^. PCR duplicates were identified and removed using the Picard package (version 2.23.4). EM-seq data was analyzed using Msuite2 software (version 2.1.0)^58,59^, an all-in-one pipeline containing quality control, read alignment, and methylation calling. Prior to alignment, the tailing 25 bp in read 1 and leading 25 bp in read 2 were trimmed to avoid overhang issues inherent to cfDNA^60^; hisat2 software (version 2.2.1)^61^ was employed as the underlying aligner and all the other parameters were kept default. EM-seq samples with an overall mapping rate below 80%, a cytosine-to-thymine conversion rate below 98%, fewer than 40 million reportable alignments, or abnormal methylation levels in promoter regions were excluded from downstream analysis. For all datasets, samples with more than 100 million reads were randomly down-sampled to 60 million reads for downstream analyses. For hepatocellular carcinoma (HCC) samples, tumor DNA fractions were calculated using ichorCNA algorithm (version 0.3.2)^62^ with a window size of 500 kb, as described in our previous study^12^.

### Identification of somatic variants in cfDNA

For both whole genome sequencing and EM-seq data, unmapped reads and secondary alignments were removed. Variant calling was then performed using the “mpileup” function in BCFtools software (version 1.12)^63^ with “-q 60” option to exclude alignments with a mapping quality lower than 60 and “-Q 30” option to exclude bases with a sequencing quality score below 30. The pileup sequences were subsequently processed with the “call” function and “-c” option in BCFtools to identify candidate variants. For EM-seq data, due to C-to-T conversion during library preparation, cytosines in the reference genome were treated differently: thymines in the sequencing data were not considered as candidate variants. Variants involving insertions/deletions, those on sex chromosomes or mitochondria, loci with more than one non-reference allele, and variants with a detection quality below 30 were all discarded.

For human samples, variants overlapping with genomic regions defined as analytically problematic by the ENCODE project^64^, as well as variants within the top 1% of coverage, were also discarded. We then filtered out variants annotated in dbSNP database (version 156)^65^ with an allele frequency greater than 0.1%. As all samples analyzed in this study were from east-Asian patients, variants with an allele frequency greater than 0.1% in the Nyuwa^66^, ChinaMAP^67^, Korea1K^68^, or ToMMo^69^ databases were also removed. Cancer-associated mutations from the COSMIC database (version 99) were retained for further analyses. For the Bie et al. dataset, one sample (accession ID: HRR1236003) was excluded due to an abnormally high number of variants. For murine cfDNA samples, variants within the top 10% of coverage were filtered out. The remaining variants, all of which were substitutions on autosomes, were retained for downstream analyses.

### Mutation profiles and deconvolution analysis

The “MutationalPatterns” package (version 3.12)^70^ was used to calculate and normalize the frequencies of 96 mutational profiles for each sample. The normalized mutation profiles were then subjected to principal component analysis (PCA) and unsupervised clustering using the “ComplexHeatmap” package (version 2.18)^71^. The 86 single-base substitution mutational signatures were downloaded from the COSMIC database (release v3.4). Signatures annotated as sequencing artifacts (N=19) were excluded, leaving 67 signatures for deconvolution.

To infer tumor-associated mutational signatures in cfDNA, a panel of control subjects (N=8, 10, and 10 for our HCC cohort, Liang et al., and Bie et al. datasets, respectively) was randomly selected to simulate the background noises in cfDNA, which includes artifacts arising from experimental or sequencing errors and clonal hematopoiesis. The mutation profiles of these controls were combined with those in COSMIC to form a reference pool for deconvolution. The non-negative matrix factorization algorithm, implemented in the “NMF” package (version 0.27)^72^, was then applied to deconvolute the mutational profiles of the remaining samples. For each cfDNA sample, the predicted contributions from the COSMIC signatures were considered “tumor-associated” and used for downstream analyses.

### Fragmentomic and epigenomic features in cfDNA

For each cfDNA sample, reads overlapping somatic variant loci identified within the sample were extracted. Reads that covered multiple variants were excluded. For all variant loci, reads carrying the reference alleles were classified as “Wt-DNA”, while those compassing variant alleles were classified as “Mut-DNA”. Fragmentomic features were then calculated separately for the Wt-DNA and Mut-DNA reads for all samples.

CfDNA molecules shorter than or equal to 150 bp (i.e., ≤ 150 bp) were considered as short fragments, and the fraction of short fragments was calculated as a size feature for each sample. For motif analysis, as described in our previous studies^10,73^, 4-mer sequences at the 5’ end of cfDNA fragments were extracted, and the frequency of each 4-mer sequence was calculated. The fraction of CCCA sequences was used to assess the 5’-CCCA end motif usage. Additionally, 4-mer sequences using 2 bases upstream and 2 bases downstream of the 5’ end were calculated, with the fraction of CTCC sequences used to assess the CT-5’-CC breakpoint motif usage^31^.

E-index values and the fraction of fragment ends located within nucleosomes were calculated as previously described^1,12^. Briefly, cfDNA ends were analyzed in an orientation-aware manner^34^: for each cfDNA molecule, its fragment ends with lower and higher genome coordinates were termed as upstream (U) and downstream (D) end, respectively, and processed separately in downstream analyses. To calculate E-index, cfDNA reads from a pool of healthy controls were collected to build a model of end distribution, and the consistency of cfDNA ends in the sample-of-interest with this model was calculated using a weighted average approach (referred to as the E-index) as illustrated in the following formula:

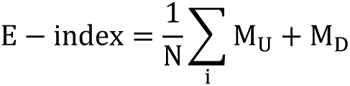

where *N* denoted the read number of the working sample (e.g., Wt-DNA), *i* denoted each sequencing read, *MU*, and *MD* denoted the counts of its ending positions serving as *U* and *D* ends in the model, respectively.

For each cfDNA end, the nearest nucleosome annotated in the GM12878 cell line (lymphoid lineage) was identified^12,32^. The distance between the cfDNA end and the nucleosome center was calculated, with negative distances assigned when the cfDNA end was upstream of the nucleosome center. The fraction of cfDNA fragment ends located within ± 50 bp of nucleosome centers was calculated using the following formula^1,12,34^:

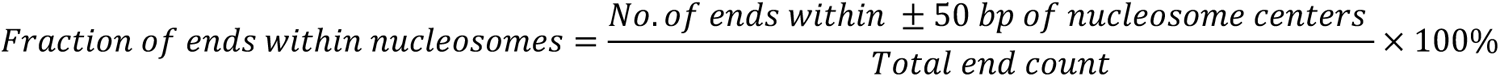

For DNA methylation analysis, only variant loci with reads covering both the reference and variant alleles were retained for comparison of the same set of CpGs. DNA methylation levels for Wt- and Mut-DNA were separately called using the “meth.caller.CpG” and “pair.CpG” programs in Msuite2 software with default parameters. For each sample, the CpG sites were divided into three groups: promoters (defined as ± 500 bp around transcription start sites), gene bodies, and intergenic regions; then DNA methylation densities were calculated for these three groups separately. DNA methylation density was calculated as the fraction of Cytosines at CpG sites in the sequencing data as shown in the following formula:

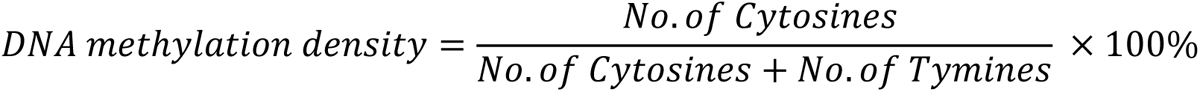

where Cytosines indicated methylated CpG while Thymies indicated unmethylated CpG in EM-seq protocol.

### Building FreeSV and FreeSV-m models

To build FreeSV model, all the somatic-variant associated features, including mutation profiles (N=96), Diff-size, Diff-CCCA, Diff-CTCC, Diff-nucleosome, and Diff-methylation for CpGs in gene bodies (totally 101 features) were used. A gradient boosted decision tree (GBDT) algorithm, implemented in the “gbm” package (version 2.1.9) in R software (version 4.2.0), was used to build diagnostic models using these features. Hyperparameter tuning was performed using the “caret” package (version 6.0.94). A 10-fold nested cross-validation was applied as follows: the samples were randomly split into 10 equal-sized subsets, with 9 subsets used for training and the remaining subset used for testing. This process was repeated for each subset, and the prediction results for each testing subset were collected. Hence, for each sample, it was predicted by a model trained using samples without itself to eliminate information disclosure. The whole procedure was repeated 100 times, and the average prediction score for each sample was calculated and determined as its prediction score. The area under the receiver operating characteristic (ROC) curve (AUC) was used to quantify model performance.

To build FreeSV-m model, besides the features used in FreeSV, four additional genomewide features (DNA methylation, fragment size, end motif, and copy number) used in THEMIS model^37^ was also leveraged (totally 105 features). For a fair comparison with THEMIS model, we adapted the data separation in Bie et al.^37^ where the samples were divided into training and testing groups. The FreeSV-m model was trained on the training group using the same approach as FreeSV; the trained model was then directly applied to the testing group. Besides ROC analyses, a specificity of 99% (which parameter was used in Bie et al.^37^) was employed to evaluate and compare the sensitives of FreeSV-m and THEMIS models.

### Statistical analysis and reproducibility

All statistical analyses were performed using R software (version 4.2.0). Parametric tests (e.g., paired t-test) were used to compare features in paired Mut- and Wt-DNA samples, while non-parametric tests (e.g., Mann-Whitney U test) were employed for other comparisons. In the Liang et al. dataset, post-treatment lung cancer samples (N=2) were excluded from the analysis. In the Bie et al. dataset, 92 samples were discarded during quality control. For all cfDNA data, samples with more than 100 million reads were randomly down sampled to 60 million reads for downstream analyses.

## Data and code availability

Raw cfDNA whole genome sequencing data for the HCC cohort were retrieved from Genome Sequence Archive in National Genomics Data Center (GSA-Human) under accession number HRA005521. Public datasets were obtained from CNGB Nucleotide Sequence Archive with accession number CNP0000680, and GSA-Human with accession number HRA003209. The nucleosome track for GM12878 cell line was downloaded from NucMap database^74^, which recorded ∼13.9 million nucleosome-protected regions accounting for ∼90% of the human genome.

## Author contributions

Conception, design, and study supervision: K.S.; Development of methodology: Z.Z., Y.A., K.S.; Acquisition of data: all authors; Analysis and interpretation of data: all authors; Writing the manuscript: Z.Z., Y.A., K.S.; Review and edit of the manuscript: K.S.

## Acknowledgements

This work was supported by National Natural Science Foundation of China (32401206), Guangdong Basic and Applied Basic Research Foundation (2023B1515120073), Shenzhen Clinical Research Center for Oral Diseases (20210617170745001), National Key R&D Program of China (2022YFA0912700), Major Program of Shenzhen Bay Laboratory (S241101004), and Shenzhen Bay Scholar Fellowship (to C.D. and K.S.). We’d like to thank Ms. Qi Wang for technical assistance, and Shenzhen Bay Laboratory HPC facility for computational support. We also thank Drs. Fenglong Bie and Shugeng Gao, Chinese Academy of Medical Sciences and Peking Union Medical College, for data sharing.

## References

1. An, Y., Zhao, X., Zhang, Z., Xia, Z., Yang, M., Ma, L., Zhao, Y., Xu, G., Du, S., Wu, X., et al. (2023). DNA methylation analysis explores the molecular basis of plasma cell-free DNA fragmentation. Nat Commun 14, 287. 10.1038/s41467-023-35959-6.

2. Sun, K., Jiang, P., Chan, K.C.A., Wong, J., Cheng, Y.K., Liang, R.H., Chan, W.K., Ma, E.S., Chan, S.L., Cheng, S.H., et al. (2015). Plasma DNA tissue mapping by genome-wide methylation sequencing for noninvasive prenatal, cancer, and transplantation assessments. Proc Natl Acad Sci U S A 112, E5503–5512. 10.1073/pnas.1508736112.

3. Lui, Y.Y., Chik, K.W., Chiu, R.W., Ho, C.Y., Lam, C.W., and Lo, Y.M. (2002). Predominant hematopoietic origin of cell-free DNA in plasma and serum after sex-mismatched bone marrow transplantation. Clin Chem 48, 421–427.

4. Moss, J., Ben-Ami, R., Shai, E., Gal-Rosenberg, O., Kalish, Y., Klochendler, A., Cann, G., Glaser, B., Arad, A., Shemer, R., and Dor, Y. (2023). Megakaryocyte- and erythroblast-specific cell-free DNA patterns in plasma and platelets reflect thrombopoiesis and erythropoiesis levels. Nat Commun 14, 7542. 10.1038/s41467-023-43310-2.

5. Stroun, M., Anker, P., Maurice, P., Lyautey, J., Lederrey, C., and Beljanski, M. (1989). Neoplastic characteristics of the DNA found in the plasma of cancer patients. Oncology 46, 318–322. 10.1159/000226740.

6. Wan, J.C.M., Massie, C., Garcia-Corbacho, J., Mouliere, F., Brenton, J.D., Caldas, C., Pacey, S., Baird, R., and Rosenfeld, N. (2017). Liquid biopsies come of age: towards implementation of circulating tumour DNA. Nat Rev Cancer 17, 223–238. 10.1038/nrc.2017.7.

7. Chan, K.C.A., Jiang, P., Chan, C.W., Sun, K., Wong, J., Hui, E.P., Chan, S.L., Chan, W.C., Hui, D.S., Ng, S.S., et al. (2013). Noninvasive detection of cancer-associated genome-wide hypomethylation and copy number aberrations by plasma DNA bisulfite sequencing. Proc Natl Acad Sci U S A 110, 18761–18768. 10.1073/pnas.1313995110.

8. Lo, Y.M.D., Han, D.S.C., Jiang, P., and Chiu, R.W.K. (2021). Epigenetics, fragmentomics, and topology of cell-free DNA in liquid biopsies. Science 372, eaaw3616. 10.1126/science.aaw3616.

9. Underhill, H.R., Kitzman, J.O., Hellwig, S., Welker, N.C., Daza, R., Baker, D.N., Gligorich, K.M., Rostomily, R.C., Bronner, M.P., and Shendure, J. (2016). Fragment length of circulating tumor DNA. PLoS Genet 12, e1006162. 10.1371/journal.pgen.1006162.

10. Jiang, P., Sun, K., Peng, W., Cheng, S.H., Ni, M., Yeung, P.C., Heung, M.M.S., Xie, T., Shang, H., Zhou, Z., et al. (2020). Plasma DNA End-Motif Profiling as a Fragmentomic Marker in Cancer, Pregnancy, and Transplantation. Cancer Discov 10, 664–673. 10.1158/2159-8290.CD-19-0622.

11. Guo, W., Chen, X., Liu, R., Liang, N., Ma, Q., Bao, H., Xu, X., Wu, X., Yang, S., Shao, Y., et al. (2022). Sensitive detection of stage I lung adenocarcinoma using plasma cell-free DNA breakpoint motif profiling. EBioMedicine 81, 104131. 10.1016/j.ebiom.2022.104131.

12. Ju, J., Zhao, X., An, Y., Yang, M., Zhang, Z., Liu, X., Hu, D., Wang, W., Pan, Y., Xia, Z., et al. (2024). Cell-free DNA end characteristics enable accurate and sensitive cancer diagnosis. Cell Rep Methods 4, 100877. 10.1016/j.crmeth.2024.100877.

13. van der Pol, Y., and Mouliere, F. (2019). Toward the early detection of cancer by decoding the epigenetic and environmental fingerprints of cell-free DNA. Cancer Cell 36, 350–368. 10.1016/j.ccell.2019.09.003.

14. Thierry, A.R. (2023). Circulating DNA fragmentomics and cancer screening. Cell Genom 3, 100242. 10.1016/j.xgen.2022.100242.

15. Liu, Y. (2022). At the dawn: cell-free DNA fragmentomics and gene regulation. Br J Cancer 126, 379–390. 10.1038/s41416-021-01635-z.

16. Zviran, A., Schulman, R.C., Shah, M., Hill, S.T.K., Deochand, S., Khamnei, C.C., Maloney, D., Patel, K., Liao, W., Widman, A.J., et al. (2020). Genome-wide cell-free DNA mutational integration enables ultra-sensitive cancer monitoring. Nat Med 26, 1114–1124. 10.1038/s41591-020-0915-3.

17. Chabon, J.J., Hamilton, E.G., Kurtz, D.M., Esfahani, M.S., Moding, E.J., Stehr, H., Schroers-Martin, J., Nabet, B.Y., Chen, B., Chaudhuri, A.A., et al. (2020). Integrating genomic features for non-invasive early lung cancer detection. Nature 580, 245–251. 10.1038/s41586-020-2140-0.

18. Jiang, P., Sun, K., Tong, Y.K., Cheng, S.H., Cheng, T.H.T., Heung, M.M.S., Wong, J., Wong, V.W.S., Chan, H.L.Y., Chan, K.C.A., et al. (2018). Preferred end coordinates and somatic variants as signatures of circulating tumor DNA associated with hepatocellular carcinoma. Proc Natl Acad Sci U S A 115, E10925–E10933. 10.1073/pnas.1814616115.

19. Li, S., Noor, Z.S., Zeng, W., Stackpole, M.L., Ni, X., Zhou, Y., Yuan, Z., Wong, W.H., Agopian, V.G., Dubinett, S.M., et al. (2021). Sensitive detection of tumor mutations from blood and its application to immunotherapy prognosis. Nat Commun 12, 4172. 10.1038/s41467-021-24457-2.

20. Chen, H., Zhang, Y., Wang, B., Liao, R., Duan, X., Yang, C., Chen, J., Hao, Y., Shu, Y., Cai, L., et al. (2024). Characterization and mitigation of artifacts derived from NGS library preparation due to structure-specific sequences in the human genome. BMC Genomics 25, 227. 10.1186/s12864-024-10157-w.

21. Liu, J., Chen, X., Wang, J., Zhou, S., Wang, C.L., Ye, M.Z., Wang, X.Y., Song, Y., Wang, Y.Q., Zhang, L.T., et al. (2019). Biological background of the genomic variations of cf-DNA in healthy individuals. Ann Oncol 30, 464–470. 10.1093/annonc/mdy513.

22. Razavi, P., Li, B.T., Brown, D.N., Jung, B., Hubbell, E., Shen, R., Abida, W., Juluru, K., De Bruijn, I., Hou, C., et al. (2019). High-intensity sequencing reveals the sources of plasma circulating cell-free DNA variants. Nat Med. 10.1038/s41591-019-0652-7.

23. Sun, K. (2019). Clonal hematopoiesis: background player in plasma cell-free DNA variants. Ann Transl Med 7, S384. 10.21037/atm.2019.12.97.

24. Sondka, Z., Dhir, N.B., Carvalho-Silva, D., Jupe, S., Madhumita, McLaren, K., Starkey, M., Ward, S., Wilding, J., Ahmed, M., et al. (2024). COSMIC: a curated database of somatic variants and clinical data for cancer. Nucleic Acids Res 52, D1210–D1217. 10.1093/nar/gkad986.

25. Alexandrov, L.B., Kim, J., Haradhvala, N.J., Huang, M.N., Tian Ng, A.W., Wu, Y., Boot, A., Covington, K.R., Gordenin, D.A., Bergstrom, E.N., et al. (2020). The repertoire of mutational signatures in human cancer. Nature 578, 94–101. 10.1038/s41586-020-1943-3.

26. Wan, J.C.M., Stephens, D., Luo, L., White, J.R., Stewart, C.M., Rousseau, B., Tsui, D.W.Y., and Diaz, L.A., Jr. (2022). Genome-wide mutational signatures in low-coverage whole genome sequencing of cell-free DNA. Nat Commun 13, 4953. 10.1038/s41467-022-32598-1.

27. Bruhm, D.C., Mathios, D., Foda, Z.H., Annapragada, A.V., Medina, J.E., Adleff, V., Chiao, E.J., Ferreira, L., Cristiano, S., White, J.R., et al. (2023). Single-molecule genome-wide mutation profiles of cell-free DNA for non-invasive detection of cancer. Nat Genet 55, 1301–1310. 10.1038/s41588-023-01446-3.

28. Lun, F.M.F., Chiu, R.W.K., Sun, K., Leung, T.Y., Jiang, P., Chan, K.C.A., Sun, H., and Lo, Y.M.D. (2013). Noninvasive prenatal methylomic analysis by genomewide bisulfite sequencing of maternal plasma DNA. Clin Chem 59, 1583–1594. 10.1373/clinchem.2013.212274.

29. Vaisvila, R., Ponnaluri, V.K.C., Sun, Z., Langhorst, B.W., Saleh, L., Guan, S., Dai, N., Campbell, M.A., Sexton, B.S., Marks, K., et al. (2021). Enzymatic methyl sequencing detects DNA methylation at single-base resolution from picograms of DNA. Genome Res 31, 1280–1289. 10.1101/gr.266551.120.

30. Schulze, K., Imbeaud, S., Letouze, E., Alexandrov, L.B., Calderaro, J., Rebouissou, S., Couchy, G., Meiller, C., Shinde, J., Soysouvanh, F., et al. (2015). Exome sequencing of hepatocellular carcinomas identifies new mutational signatures and potential therapeutic targets. Nat Genet 47, 505–511. 10.1038/ng.3252.

31. Jin, X., Wang, Y.Q., Xu, J.J., Li, Y.M., Cheng, F.J., Luo, Y.X., Zhou, H.B., Lin, S.W., Xiao, F., Zhang, L., et al. (2023). Plasma cell-free DNA promise monitoring and tissue injury assessment of COVID-19. Molecular Genetics and Genomics 298, 823–836. 10.1007/s00438-023-02014-4.

32. Snyder, M.W., Kircher, M., Hill, A.J., Daza, R.M., and Shendure, J. (2016). Cell-free DNA comprises an in vivo nucleosome footprint that informs its tissues-of-origin. Cell 164, 57–68. 10.1016/j.cell.2015.11.050.

33. Rao, S., Han, A.L., Zukowski, A., Kopin, E., Sartorius, C.A., Kabos, P., and Ramachandran, S. (2022). Transcription factor-nucleosome dynamics from plasma cfDNA identifies ER-driven states in breast cancer. Sci Adv 8, eabm4358. 10.1126/sciadv.abm4358.

34. Sun, K., Jiang, P., Cheng, S.H., Cheng, T.H.T., Wong, J., Wong, V.W.S., Ng, S.S.M., Ma, B.B.Y., Leung, T.Y., Chan, S.L., et al. (2019). Orientation-aware plasma cell-free DNA fragmentation analysis in open chromatin regions informs tissue of origin. Genome Res 29, 418–427. 10.1101/gr.242719.118.

35. Sarsani, V.K., Raghupathy, N., Fiddes, I.T., Armstrong, J., Thibaud-Nissen, F., Zinder, O., Bolisetty, M., Howe, K., Hinerfeld, D., Ruan, X., et al. (2019). The Genome of C57BL/6J “Eve”, the Mother of the Laboratory Mouse Genome Reference Strain. G3 (Bethesda) 9, 1795–1805. 10.1534/g3.119.400071.

36. Liang, H., Li, F., Qiao, S., Zhou, X., Xie, G., Zhao, X., Zhang, Y., and Wu, K. (2020). Whole-genome sequencing of cell-free DNA yields genome-wide read distribution patterns to track tissue of origin in cancer patients. Clin Transl Med 10, e177. 10.1002/ctm2.177.

37. Bie, F., Wang, Z., Li, Y., Guo, W., Hong, Y., Han, T., Lv, F., Yang, S., Li, S., Li, X., et al. (2023). Multimodal analysis of cell-free DNA whole-methylome sequencing for cancer detection and localization. Nat Commun 14, 6042. 10.1038/s41467-023-41774-w.

38. Zhou, X., Cheng, Z., Dong, M., Liu, Q., Yang, W., Liu, M., Tian, J., and Cheng, W. (2022). Tumor fractions deciphered from circulating cell-free DNA methylation for cancer early diagnosis. Nat Commun 13, 7694. 10.1038/s41467-022-35320-3.

39. Ambrosi, C., Manzo, M., and Baubec, T. (2017). Dynamics and Context-Dependent Roles of DNA Methylation. J Mol Biol 429, 1459–1475. 10.1016/j.jmb.2017.02.008.

40. Cristiano, S., Leal, A., Phallen, J., Fiksel, J., Adleff, V., Bruhm, D.C., Jensen, S.O., Medina, J.E., Hruban, C., White, J.R., et al. (2019). Genome-wide cell-free DNA fragmentation in patients with cancer. Nature 570, 385–389. 10.1038/s41586-019-1272-6.

41. Liu, X., Yang, M., Hu, D., An, Y., Wang, W., Lin, H., Pan, Y., Ju, J., and Sun, K. (2024). Systematic biases in reference-based plasma cell-free DNA fragmentomic profiling. Cell Rep Methods 4, 100793. 10.1016/j.crmeth.2024.100793.

42. Cobo, I., Tanaka, T., Glass, C.K., and Yeang, C. (2022). Clonal hematopoiesis driven by DNMT3A and TET2 mutations: role in monocyte and macrophage biology and atherosclerotic cardiovascular disease. Curr Opin Hematol 29, 1–7. 10.1097/MOH.0000000000000688.

43. Buscarlet, M., Provost, S., Zada, Y.F., Barhdadi, A., Bourgoin, V., Lepine, G., Mollica, L., Szuber, N., Dube, M.P., and Busque, L. (2017). DNMT3A and TET2 dominate clonal hematopoiesis and demonstrate benign phenotypes and different genetic predispositions. Blood 130, 753–762. 10.1182/blood-2017-04-777029.

44. Izzo, F., Lee, S.C., Poran, A., Chaligne, R., Gaiti, F., Gross, B., Murali, R.R., Deochand, S.D., Ang, C., Jones, P.W., et al. (2020). DNA methylation disruption reshapes the hematopoietic differentiation landscape. Nat Genet 52, 378–387. 10.1038/s41588-020-0595-4.

45. McClatchy, J., Strogantsev, R., Wolfe, E., Lin, H.Y., Mohammadhosseini, M., Davis, B.A., Eden, C., Goldman, D., Fleming, W.H., Conley, P., et al. (2023). Clonal hematopoiesis related TET2 loss-of-function impedes IL1beta-mediated epigenetic reprogramming in hematopoietic stem and progenitor cells. Nat Commun 14, 8102. 10.1038/s41467-023-43697-y.

46. Teo, Y.V., Capri, M., Morsiani, C., Pizza, G., Faria, A.M.C., Franceschi, C., and Neretti, N. (2019). Cell-free DNA as a biomarker of aging. Aging Cell 18, e12890. 10.1111/acel.12890.

47. Diamandis, E.P. (2016). A Word of Caution on New and Revolutionary Diagnostic Tests. Cancer Cell 29, 141–142. 10.1016/j.ccell.2016.01.003.

48. Hollizeck, S., Wang, N., Wong, S.Q., Litchfield, C., Guinto, J., Ftouni, S., Rebello, R., Kanwal, S., Dong, R., Grimmond, S., et al. (2024). Unravelling mutational signatures with plasma circulating tumour DNA. Nat Commun 15, 9876. 10.1038/s41467-024-54193-2.

49. Pardali, E., Dimmeler, S., Zeiher, A.M., and Rieger, M.A. (2020). Clonal hematopoiesis, aging, and cardiovascular diseases. Exp Hematol 83, 95–104. 10.1016/j.exphem.2019.12.006.

50. Bolton, K.L., Ptashkin, R.N., Gao, T., Braunstein, L., Devlin, S.M., Kelly, D., Patel, M., Berthon, A., Syed, A., Yabe, M., et al. (2020). Cancer therapy shapes the fitness landscape of clonal hematopoiesis. Nat Genet 52, 1219–1226. 10.1038/s41588-020-00710-0.

51. Fairchild, L., Whalen, J., D’Aco, K., Wu, J., Gustafson, C.B., Solovieff, N., Su, F., Leary, R.J., Campbell, C.D., and Balbin, O.A. (2023). Clonal hematopoiesis detection in patients with cancer using cell-free DNA sequencing. Sci Transl Med 15, eabm8729. 10.1126/scitranslmed.abm8729.

52. Wong, D., Tageldein, M., Luo, P., Ensminger, E., Bruce, J., Oldfield, L., Gong, H., Fischer, N.W., Laverty, B., Subasri, V., et al. (2024). Cell-free DNA from germline TP53 mutation carriers reflect cancer-like fragmentation patterns. Nat Commun 15, 7386. 10.1038/s41467-024-51529-w.

53. Olova, N., Krueger, F., Andrews, S., Oxley, D., Berrens, R.V., Branco, M.R., and Reik, W. (2018). Comparison of whole-genome bisulfite sequencing library preparation strategies identifies sources of biases affecting DNA methylation data. Genome Biol 19, 33. 10.1186/s13059-018-1408-2.

54. Meddeb, R., Pisareva, E., and Thierry, A.R. (2019). Guidelines for the Preanalytical Conditions for Analyzing Circulating Cell-Free DNA. Clin Chem 65, 623–633. 10.1373/clinchem.2018.298323.

55. Sun, K. (2020). Ktrim: an extra-fast and accurate adapter- and quality-trimmer for sequencing data. Bioinformatics 36, 3561–3562. 10.1093/bioinformatics/btaa171.

56. Li, H., and Durbin, R. (2009). Fast and accurate short read alignment with Burrows-Wheeler transform. Bioinformatics 25, 1754–1760. 10.1093/bioinformatics/btp324.

57. Li, H., Handsaker, B., Wysoker, A., Fennell, T., Ruan, J., Homer, N., Marth, G., Abecasis, G., Durbin, R., and Genome Project Data Processing, S. (2009). The Sequence Alignment/Map format and SAMtools. Bioinformatics 25, 2078–2079. 10.1093/bioinformatics/btp352.

58. Li, L., An, Y., Ma, L., Yang, M., Yuan, P., Liu, X., Jin, X., Zhao, Y., Zhang, S., Hong, X., and Sun, K. (2022). Msuite2: All-in-one DNA methylation data analysis toolkit with enhanced usability and performance. Comput Struct Biotechnol J 20, 1271–1276. 10.1016/j.csbj.2022.03.005.

59. Sun, K., Li, L., Ma, L., Zhao, Y., Deng, L., Wang, H., and Sun, H. (2020). Msuite: A High-Performance and Versatile DNA Methylation Data-Analysis Toolkit. Patterns (N Y) 1, 100127. 10.1016/j.patter.2020.100127.

60. Wu, T., Chen, W., Yang, Z., Tan, H., Wang, J., Xiao, X., Li, M., and Zhao, M. (2018). DNA terminal structure-mediated enzymatic reaction for ultra-sensitive discrimination of single nucleotide variations in circulating cell-free DNA. Nucleic Acids Res 46, e24. 10.1093/nar/gkx1218.

61. Kim, D., Paggi, J.M., Park, C., Bennett, C., and Salzberg, S.L. (2019). Graph-based genome alignment and genotyping with HISAT2 and HISAT-genotype. Nat Biotechnol 37, 907–915. 10.1038/s41587-019-0201-4.

62. Adalsteinsson, V.A., Ha, G., Freeman, S.S., Choudhury, A.D., Stover, D.G., Parsons, H.A., Gydush, G., Reed, S.C., Rotem, D., Rhoades, J., et al. (2017). Scalable whole-exome sequencing of cell-free DNA reveals high concordance with metastatic tumors. Nat Commun 8, 1324. 10.1038/s41467-017-00965-y.

63. Danecek, P., Bonfield, J.K., Liddle, J., Marshall, J., Ohan, V., Pollard, M.O., Whitwham, A., Keane, T., McCarthy, S.A., Davies, R.M., and Li, H. (2021). Twelve years of SAMtools and BCFtools. Gigascience 10. 10.1093/gigascience/giab008.

64. Amemiya, H.M., Kundaje, A., and Boyle, A.P. (2019). The ENCODE Blacklist: Identification of Problematic Regions of the Genome. Sci Rep 9, 9354. 10.1038/s41598-019-45839-z.

65. Sherry, S.T., Ward, M.H., Kholodov, M., Baker, J., Phan, L., Smigielski, E.M., and Sirotkin, K. (2001). dbSNP: the NCBI database of genetic variation. Nucleic Acids Res 29, 308–311. 10.1093/nar/29.1.308.

66. Zhang, P., Luo, H., Li, Y., Wang, Y., Wang, J., Zheng, Y., Niu, Y., Shi, Y., Zhou, H., Song, T., et al. (2021). NyuWa Genome resource: A deep whole-genome sequencing-based variation profile and reference panel for the Chinese population. Cell Rep 37, 110017. 10.1016/j.celrep.2021.110017.

67. Cao, Y., Li, L., Xu, M., Feng, Z., Sun, X., Lu, J., Xu, Y., Du, P., Wang, T., Hu, R., et al. (2020). The ChinaMAP analytics of deep whole genome sequences in 10,588 individuals. Cell Res 30, 717–731. 10.1038/s41422-020-0322-9.

68. Jeon, S., Bhak, Y., Choi, Y., Jeon, Y., Kim, S., Jang, J., Jang, J., Blazyte, A., Kim, C., Kim, Y., et al. (2020). Korean Genome Project: 1094 Korean personal genomes with clinical information. Sci Adv 6, eaaz7835. 10.1126/sciadv.aaz7835.

69. Koshiba, S., Motoike, I., Saigusa, D., Inoue, J., Shirota, M., Katoh, Y., Katsuoka, F., Danjoh, I., Hozawa, A., Kuriyama, S., et al. (2018). Omics research project on prospective cohort studies from the Tohoku Medical Megabank Project. Genes Cells 23, 406–417. 10.1111/gtc.12588.

70. Manders, F., Brandsma, A.M., de Kanter, J., Verheul, M., Oka, R., van Roosmalen, M.J., van der Roest, B., van Hoeck, A., Cuppen, E., and van Boxtel, R. (2022). MutationalPatterns: the one stop shop for the analysis of mutational processes. BMC Genomics 23, 134. 10.1186/s12864-022-08357-3.

71. Gu, Z., Eils, R., and Schlesner, M. (2016). Complex heatmaps reveal patterns and correlations in multidimensional genomic data. Bioinformatics 32, 2847–2849. 10.1093/bioinformatics/btw313.

72. Gaujoux, R., and Seoighe, C. (2010). A flexible R package for nonnegative matrix factorization. BMC Bioinformatics 11, 367. 10.1186/1471-2105-11-367.

73. Serpas, L., Chan, R.W.Y., Jiang, P., Ni, M., Sun, K., Rashidfarrokhi, A., Soni, C., Sisirak, V., Lee, W.S., Cheng, S.H., et al. (2019). Dnase1l3 deletion causes aberrations in length and end-motif frequencies in plasma DNA. Proc Natl Acad Sci U S A 116, 641–649. 10.1073/pnas.1815031116.

74. Zhao, Y., Wang, J., Liang, F., Liu, Y., Wang, Q., Zhang, H., Jiang, M., Zhang, Z., Zhao, W., Bao, Y., et al. (2019). NucMap: a database of genome-wide nucleosome positioning map across species. Nucleic Acids Res 47, D163–D169. 10.1093/nar/gky980.

